# Amyloid structure determination in RELION-3.1

**DOI:** 10.1101/823310

**Authors:** Sjors H.W. Scheres

## Abstract

Helical reconstruction in RELION is increasingly used to determine atomic structures of amyloid filaments from electron cryo-microscopy (cryo-EM) images. However, because the energy landscape of amyloid refinements is typically fraught with local optima, amyloid structure determination is often difficult. This paper aims to help RELION users in this process. It discusses aspects of helical reconstruction that are specific to amyloids; it illustrates the problem of local optima in refinement and how to detect these; and it introduces a new method to calculate 3D initial models from reference-free 2D class averages. By providing starting models that are closer to the global optimum, this method makes amyloid structure determination easier. All methods described are open-source and distributed within RELION-3.1. Their use is illustrated using a publicly available data set on tau filaments from the brain of an individual with Alzheimer’s disease.

## 1. Introduction

In some aspects, cryo-EM structure determination of objects with helical symmetry is easier than single-particle analysis of globular objects. Provided the helix is long enough, a single projection image of a helix contains all necessary views for three-dimensional (3D) reconstruction of the structure, up to a limited resolution. Moreover, if the helical symmetry is known, all these views have known relative orientations. It is therefore perhaps not surprising that the first 3D reconstruction of a biological object from electron microscopy images was that of a helical object: the extended tail of the T4 bacteriophage (De Rosier & Klug, 1968; DeRosier & Moore, 1970).

Initially, EM structures of helical objects were solved by Fourier-Bessel inversion (Cochran *et al.*, 1952; Klug *et al.*, 1958). This technique requires near-perfect helical symmetry in the sample and many consider it to be difficult to use (see (Diaz *et al.*, 2010) for a review). An alternative method to solve structures of helical objects is analogous to the single-particle approach for globular objects. In this method, one divides images of individual helices into multiple smaller segments, which are boxed out individually and aligned against a common 3D reference structure by projection matching. The introduction of many more alignment parameters allows modelling deviations from helical symmetry.

Early applications of the single-particle-like approach to helical reconstruction were done in real-space (Bluemke *et al.*, 1988; Sosa *et al.*, 1997; Beroukhim & Unwin, 1997). The real-space approach was opened up for wide application by implementation of the iterative helical real-space reconstruction, or IHRSR, algorithm (Egelman, 2000). More recently, we implemented a similar approach in the empirical Bayesian framework of RELION, which operates in Fourier-space (He & Scheres, 2017). The statistical framework allows expressing expectations about deviations from helical symmetry through Gaussian-shaped priors on the relative orientations of the individual segments. We used this implementation to solve the first cryo-EM structures of amyloid filaments to sufficient resolution to build *de novo* atomic models: the paired helical and straight filaments (PHFs and SFs) that are formed by the tau protein in Alzheimer’s disease (Fitzpatrick *et al.*, 2017). Since then, the approach in RELION has been used to solve multiple cryo-EM structures of amyloids, e.g. from tau (Falcon *et al.*, 2018*a*; Falcon *et al.*, 2018*b*; Falcon *et al.*, 2019; Zhang *et al.*, 2019), *α*-synuclein (Li *et al.*, 2018*b*; Li *et al.*, 2018*a*; Guerrero-Ferreira *et al.*, 2018; Guerrero-Ferreira *et al.*, 2019; Boyer *et al.*, 2019), *β*2-microglobulin (Iadanza *et al.*, 2018), amyloid protein A and amyloid light-chain from systemic amyloidosis (Liberta *et al.*, 2019; Swuec *et al.*, 2019; Radamaker *et al.*, 2019), TDP-43 (Cao *et al.*, 2019) and amyloid*-β* (Kollmer *et al.*, 2019).

Amyloid filaments are helical aggregates of proteins that are characterised by a cross-*β*-sheet quaternary structure. A few dozen proteins in the human genome are known to form amyloids, associated with more than 50 human diseases (Knowles *et al.*, 2014). In the cross-*β* arrangement, *β*-sheets stack along the direction of the helical axis and pack against each other in the plane perpendicular to the helical axis. The absence of larger structural features along the helical axis typically complicates cryo-EM structure determination of amyloids to high resolution. This is because alignment of individual segments along the helical axis depends solely on the signal caused by the 4.75 Å distance between adjacent *β*-strands.

Whereas imposing incorrect helical symmetry is a well-described pitfall for helical reconstruction (Egelman, 2000; Egelman, 2014), determining the helical symmetry for amyloids is often relatively straightforward. In RELION, and many other single-particle based programs, the helical symmetry is expressed by two parameters: the helical rise and the helical twist. Provided the filaments show discernable cross-overs in the raw micrographs or in 2D class averages, the twist can simply be calculated from the knowledge that the filament twists 180° in a single cross-over distance (*d* in Å), such that:

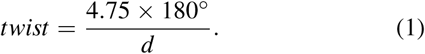

If each rung in the *β*-sheet consists of a single copy of the protein, then the rise is equal to the distance between two strands in a *β*-sheet: ∼4.75 Å. In this case, the power spectrum of micrographs with long and straight helices will show a single layer line at 4.75 Å. If a single copy of the protein folds back on itself in *m* rungs, additional layer lines at *m* × 4.75 Å will be visible and the rise will be *m* × 4.75 Å. An example of such an amyloid is the prion domain of the fungal prion HET-s, for which *m* = 2 as determined by solid-state nuclear magnetic resonance (NMR) (Van Melckebeke *et al.*, 2010). In practice, errors in the nominal magnification of cryo-EM images may be up to several percent, which leads to equally large errors in the rise. One useful way to detect nominal magnification errors is by examining the frequency of the ∼ 4.75 Å peak in the power spectrum of high-resolution 2D class averages (or in the `rlnSsnrMap` columns of the `data model class x` tables of all classes in a ‘2D classification’ job). Deviations from the expected frequency of 4.75 Å may be used to calibrate the magnification.

After an initial refinement with these estimates for the helical rise and twist, the refined map may reveal the presence of additional symmetry. Two types of additional symmetry may be present. Firstly, the amyloid may adopt *n*-fold rotational symmetry. This happens when *n* copies of the protein are rotated with respect to each other by 360/*n*°, and these *n* copies are all positioned at the same height along the helical axis. In this case, rise and twist remain as described above, and the structure is refined in symmetry point group *Cn*. An example of a C2 amyloid structure is the in-vitro reconstituted *β*2-microglobulin fibril (Iadanza et al, 2018). Alternatively, the *n* copies may be positioned at different heights along the helical axis, in a spiral staircase-like arrangement with a pseudo *n*_1_ screw axis. In this case, the structure is refined in symmetry point group *C*1, and twist and rise are calculated as described in section 2.1 of (He & Scheres, 2017). The PHFs of tau are an example with a pseudo 2_1_ screw axis; the SFs of tau are an example without additional symmetry (Fitzpatrick *et al.*, 2017).

Unfortunately, despite the relative ease with which helical symmetry parameters may be determined, cryo-EM structure determination of amyloid filaments is often far from straight-forward. This is caused by the observation that amyloid refinements are typically fraught with local optima. This characteristic is poorly described in the literature, as structures in the published literature are typically deemed to have converged onto a useful solution. Here, I illustrate the problem of amyloid refinement in RELION getting stuck in a local optimum, I describe how this problem may be detected, and I propose a program that seeks to alleviate this problem by calculating initial 3D models *de novo* from 2D reference-free class averages. The latter has already proven useful for several of our structure determination projects (Falcon *et al.*, 2018*a*; Falcon *et al.*, 2018*b*; Falcon *et al.*, 2019; Zhang *et al.*, 2019).

## 2. Methods

Reference-free 2D class averaging is a powerful method to increase signal-to-noise ratios in cryo-EM projection images without the use of any prior knowledge about the underlying structure. In RELION, the user can choose the box size of the extracted amyloid segments that are to be used for 2D class averaging or 3D refinement. Smaller boxes typically lead to higher-resolution 2D class averages, as deviations from and variations in helical symmetry within the data set have smaller effects over shorter distances.

Nevertheless, despite their lower resolution, 2D class averages that are large enough to span an entire cross-over are a convenient source of information to obtain an initial 3D model for helical objects. For reasons of computational speed, the method proposed assumes that the underlying 3D structure is devoid of any features along the helical (Z) axis, *i.e.* it consists of a single 2D structure in the XY-plane, and this structure slowly rotates in subsequent Z-planes. This assumption holds well for amyloid structures with resolutions lower than 5 Å, where the *β*-strands along the helical axis are not discernable. Subsequent columns of pixels in (horizontally aligned) 2D class averages that span an entire cross-over are then 1D projections of the underlying 2D structure in the XY-plane (Figure 1). As the relative angles of each of these 1D projections can be calculated from the observation that the entire cross-over corresponds to a 180° rotation, one can directly perform a 2D reconstruction from the collection of 1D pixel columns. The 2D reconstruction can then be inserted many times into a 3D map by rotating each Z-slice accordingly. This functionality has been implemented in the command-line program `relion helix inimodel2d` of RELION-3.1.

**Figure 1.**
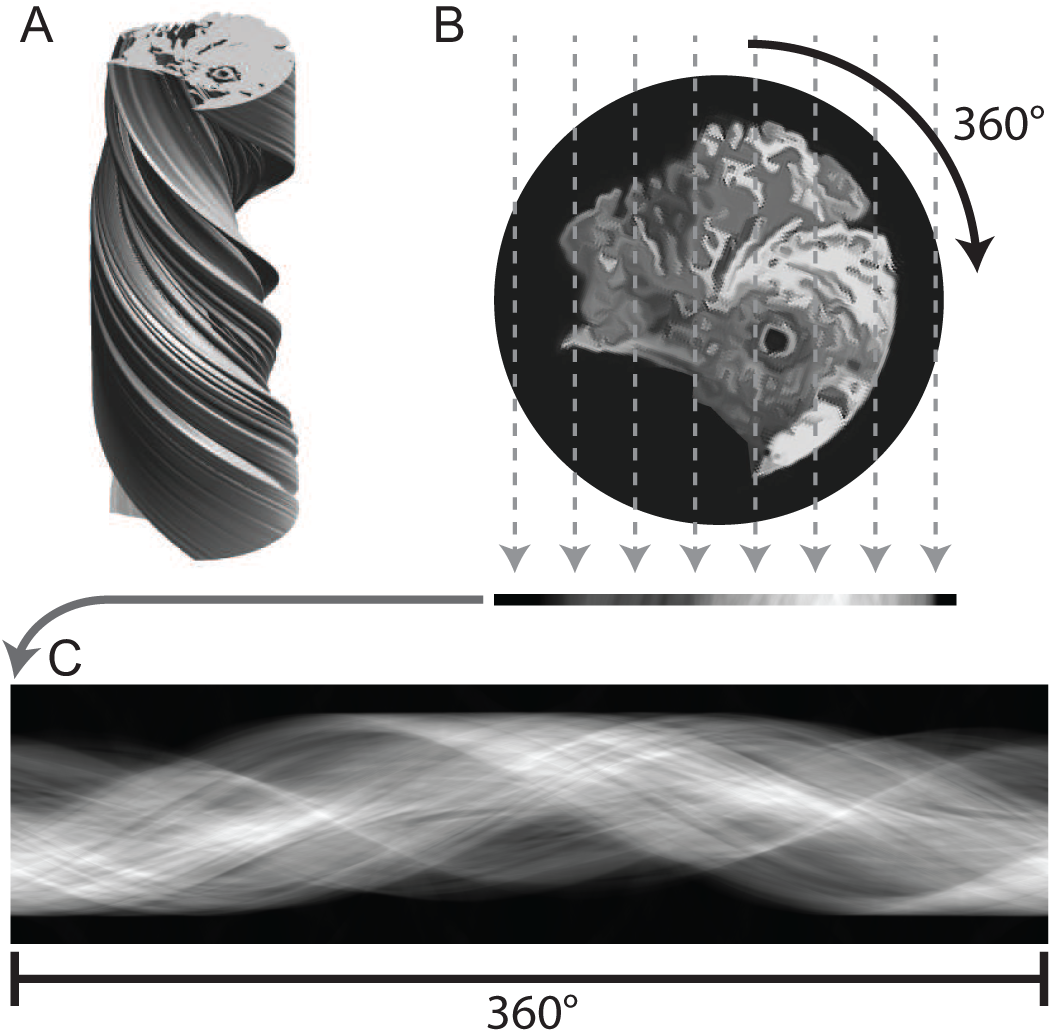
Concept of the initial model generation procedure. **A.** A 3D helical structure consists of the image of a chicken in the XY-direction and is devoid of features in the Z direction. **B.** The XY cross-section of the helix shown in A. Grey dashed lines represent a 1D projection operation, which results in a single array of pixels at the bottom of the panel. **C.** This array of pixels is inserted as a single column in the image. Repeating the 1D projection operation for 360 degrees yields a so-called sinogram image, which is shown. The initial model generation procedure outlined in the main text basically performs the inverse of this procedure: reference-free 2D class averages are aligned and summed together to form segments of a complete sinogram image, and a 2D reconstruction is performed with all pixel columns. The resulting 2D image is then inserted as multiple XY-slices with varying rotations in the 3D map.

The resolution of the resulting map is limited by the resolution of the 2D class average used. As observed above, 2D class averaging with smaller boxes typically leads to higher resolutions. Therefore, the low-resolution model obtained from a single large class average can be improved by subsequently using smaller, higher-resolution class averages. However, because multiple smaller class averages are required to cover an entire cross-over, one then also needs to determine their relative positions along the helical axis. Moreover, it is often more difficult to horizontally align smaller 2D class averages, which compounds the alignment problem further with an in-plane rotation and a translation in the Y-direction for each class average.

To this purpose, the `relion helix inimodel2d` program can also be run in an iterative manner. It takes a STAR file with all 2D class averages as input (for example as obtained from a selection of suitable classes in a ‘Subset selection’ job), together with the pixel size, the cross-over distance, and information about the step size and search range of the Y-translations and the in-plane rotations. Optionally, the user can provide an initial 2D reference image using the `--iniref` option. Provided it is re-scaled to the same pixel size and re-windowed to the same box size as the input 2D class averages (using the `relion image handler` program), this initial reference could for example come from the low-resolution XY-structure obtained with the 2D class average spanning an entire cross-over. If no initial 2D reference is provided, the program will generate one by assigning random X-positions to all 2D class averages, and performing the corresponding reconstruction from all 1D pixel columns. In an iterative manner, the 2D reference structure is then projected into 1D pixel columns that span an entire cross-over; and each of the 2D class averages is aligned with respect to this cross-over image. At the end of each iteration, a new 2D reference is then reconstructed from all corresponding pixel columns. The number of iterations is controlled by the user, and after the last iteration a 3D map is generated by rotating the final 2D reconstruction for each Z-slice according to the helical twist. Section 3 provides two examples of how to use the `relion helix inimodel2d` program. A full list of options and their explanation can be obtained from the command line by executing the program without providing any arguments.

## 3. Results

### 3.1. Test data set

The approach outlined above was tested using the motion-corrected micrograph images of EMPIAR-10230 (Iudin *et al.*, 2016), which is a cryo-EM data set on tau filaments that were extracted from the brain of an individual with sporadic Alzheimer’s disease (Falcon *et al.*, 2018*b*). These data were collected on a Gatan K2-Summit detector on a Titan Krios microscope, which was operated at an accelerating voltage of 300 kV. Inelastically scattered electrons were removed using a Gatan energy filter, using a slit width of 20 eV.

Contrast transfer function (CTF) parameters were estimated from the micrographs using the open-source software CTFFIND-4.1 (Rohou & Grigorieff, 2015). The manually picked coordinates for the paired helical filaments (PHFs) that are distributed with the EMPIAR entry were used for the extraction of individual segment images, or particles from now on. A first set of 154,643 particles were extracted using an inter-box distance of 14.1 Å and a box size of 768 pixels. Down-scaling to a box size of 256 pixels reduced the pixel size from the original 1.15 Å in the micrographs to 3.45 Å in the particles. A second set of 198,021 particles were extracted using the same inter-box distance in a box size of 256 pixels without down-scaling. Reference-free 2D class averaging was performed for both sets of particles, using 200 classes; a regularization parameter of *T* = 2; an in-plane angular sampling rate of 2°; a tube diameter of 200 Å; and a restriction of the translational offsets along the helical axis of 4.75 Å, i.e. one helical rise. The latter is a new option for 2D class averaging of helices in RELION-3.1.

### 3.2. Initial model from a single large class average

A single class average spanning an entire cross-over was selected from the 2D class averaging job with the first set of particles. This image was manually rotated and translated to align its apparent central mirror axes, an early indication of the presence of two protofilaments in the 3D structure, with the central X-axis of the image (Figure 2A). The cross-over distance was measured from the class average image to be ∼800 Å. An initial 3D model was then generated using the approach described above (Figure 2B-C). This calculation took approximately five seconds. The output map was then re-scaled and re-windowed to match the particles in the second set. The following commands were executed.

~~~
relion_image_handler
--i 180@Class2D/job051/run_it025_classes.mrcs
--o bigclass.mrc

relion_project
--i bigclass.mrc --o bigclass_ali.mrc
--psi -0.67

relion_image_handler
--i bigclass_ali.mrc --o bigclass_ali.mrc
--shift_y -5

relion_star_loopheader
rlnReferenceImage > bigclass.star

cat bigclass_ali.mrc >> bigclass.star

relion_helix_inimodel2d
--i bigclass.star --iter 1 --sym 2
--crossover_distance 800 --angpix 3.45
--o IniModel/bigclass --mask_diameter 230

relion_image_handler
--i IniModel/bigclass_class001_rec3d.mrc
--o IniModel/bigclass_class001_rec3d_box256.mrc
--angpix 3.45 --rescale_angpix 1.15 --new_box 256
~~~

**Figure 2.**
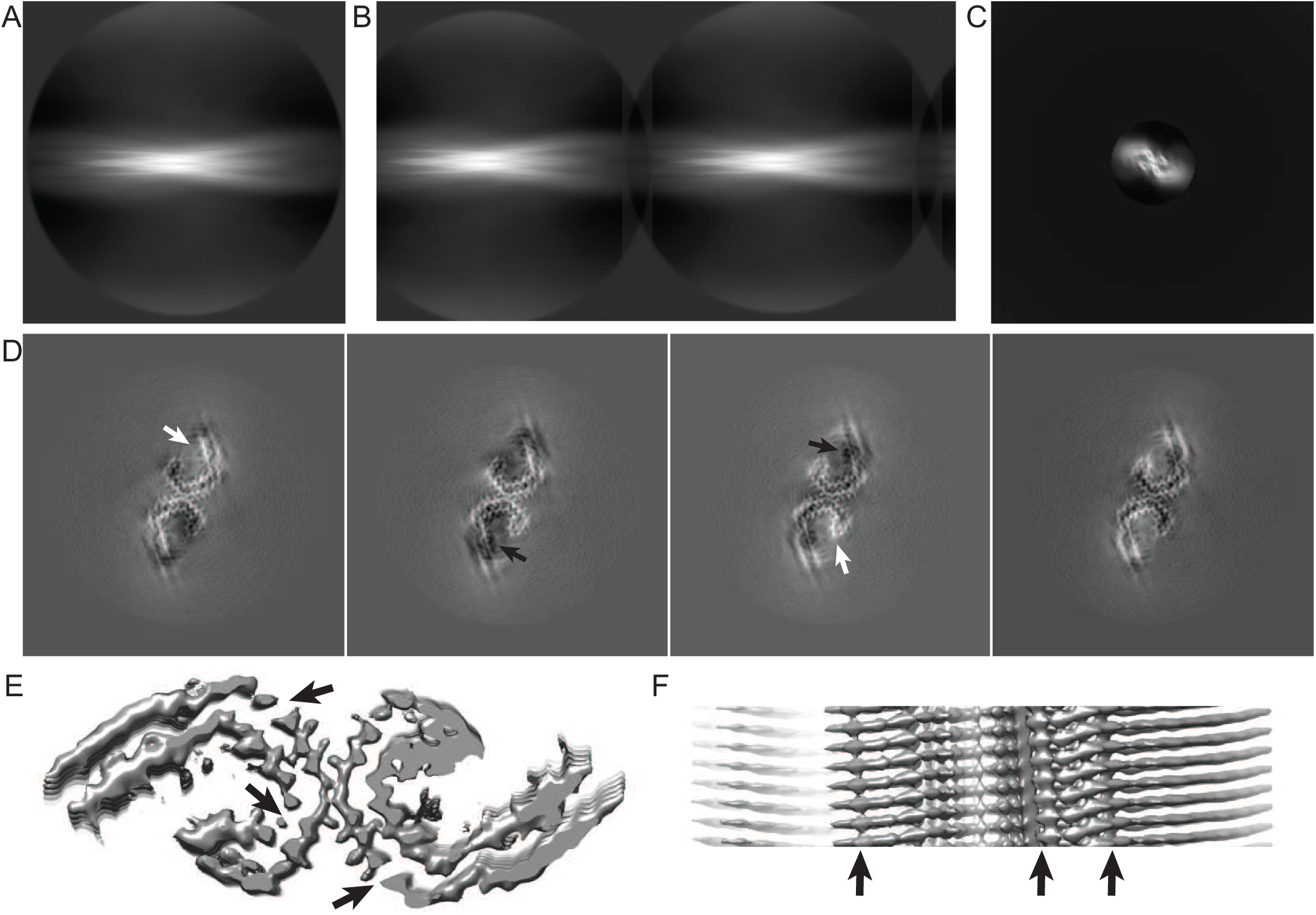
Initial model generation from a single 2D class average that spanned an entire cross-over. **A.** The 2D class average used for initial model generation. **B.** The sinogram covering 360 degrees of rotation along the helical axis, as output by RELION. Note that a single cross-over spans 180 degrees. Therefore, the image in A appears twice: one as is and once mirrored in the horizontal axis. **C.** The 2D reconstruction obtained from (half of) the pixel columns in B. **D.** Four central slices of the symmetrised, post-processing map after 3D auto-refinement, which used the initial model that was generated from the image in C. Too white areas are highlighted with white arrows; too black areas with black arrows. **E.** Thresholded view of the symmetrised, post-processing map in a view along the helical axis. Unexpected breaks in the density are highlighted with arrows. **F.** Thresholded view of the symmetrised, post-processing map in a view perpendicular to the helical axis. Unexpected connections between rungs along the helical axis are highlighted with arrows.

The resulting 3D map was then subjected to standard 3D auto-refinement with helical symmetry, using an initial resolution limit of 10 Å (through the option ‘initial low-pass filter’ on the graphical interface); C1 symmetry; an initial angular sampling of 3.7°; an initial offset search range of 8 pixels; a helical twist of −1.07°; and a helical rise of 4.75 Å. Post-processing of the refined unfiltered half-maps, using a soft mask spanning 30% of the box along the helical axis, resulted in a resolution estimate of 3.8 Å. After imposing helical symmetry in real-space, the post-processed map showed a two-fold symmetric structure reminiscent of the two C-shaped protofilaments previously observed for PHFs (Fitzpatrick *et al.*, 2017; Falcon *et al.*, 2018*b*), with clear 4.75Å separated densities along the helical axis (Figure 2D).

However, despite the reasonable resolution and the appearance of protein-like features, there are also signs that this map may represent a local optimum of refinement. When displayed at a threshold of five standard deviations, the map shows breaks in what should correspond to the main-chain density in the XY-plane (arrows in Figure 2E), while at that same threshold level densities are connected along the helical axis (arrows in Figure 2F). Moreover, the main-chain density is much stronger in some parts of the structure than in other parts, e.g. at the tip and at the ends of the C-shape protofilaments (white arrows in Figure 2D), while densities in between the 4.75 Å repeats along the helical axis are even weaker than the density in the surrounding solvent, i.e. they have negative density (black arrows in Figure 2D).

### 3.3. Initial model from multiple smaller class averages

To generate an improved initial model, 24 reference-free 2D class averages from the second set of particles were selected. To reduce the effects of the circular mask, the selected class averages were re-windowed to boxes of 220 pixels. Iterative alignment of the 24 selected, re-boxed 2D class averages of the second particle set, starting from random positions along the cross-over, took approximately 6 minutes and yielded a 3D map that was then re-windowed to a boxsize of 256 pixels. Using a cross-over distance of 700 Å resulted in better alignment of the 2D class average images along the helical axis, as compared to the 800 Å distance used in section 3.2. The following commands were executed:

~~~
relion_stack_create
--i Select/job060/class_averages.star
--o IniModel/smallclasses_box256 --ignore_optics

relion_image_handler
--i IniModel/smallclasses_box256.mrcs --new_box 220
--o IniModel/smallclasses_box220.mrcs

relion_helix_inimodel2d
--i IniModel/smallclasses_box220.star
--iter 10 --crossover_distance 700 --angpix 1.15
--o IniModel/smallclasses --mask_diameter 200
--search_shift 3 --search_angle 1 --step_angle 1
--sym 2 --maxres 5 --j 12

relion_image_handler
--i IniModel/smallclasses_class001_rec3d.mrc
--o IniModel/smallclasses_class001_rec3d_box256.mrc
--new_box 256
~~~

The resulting map was subjected to 3D auto-refinement, using an initial resolution limit of 10 Å; C1 symmetry; an initial angular sampling of 3.7°; an initial offset search range of 8 pixels; a helical twist of −1.2° and a helical rise of 4.75 Å. During this refinement, the two independently refined half-maps showed clear separation of the *β*-strands, but they converged to a different relative position along the Z-axis, resulting in near-zero FSC-values at 4.75 Å. This was remediated by aligning the two half-maps in UCSF Chimera (Pettersen *et al.*, 2004) and using the sum of the aligned half-maps as initial model for a second refinement. The second refinement used an initial resolution limit of 4.5 Å; an initial angular sampling of 1.8°; an initial offset search range of 5 pixels; and a soft-edged solvent mask with solvent-corrected FSC values. In addition, a helical twist of 179.4° and a helical rise of 2.375 Å were used to impose a pseudo 2_1_ helical screw symmetry, and these values were allowed to change during the refinement; a similar refinement with C2 symmetry gave worse results (not shown). After post-processing and imposing pseudo 2_1_ helical symmetry in real-space, the final map, with an estimated resolution of 3.2 Å, showed continuous density for the main-chain of individual monomers in all three directions. Moreover, the map showed improved side chain densities compared to the map obtained in section 3.2, and the densities for individual monomers were well-separated in the Z-direction (Figure 3).

**Figure 3.**
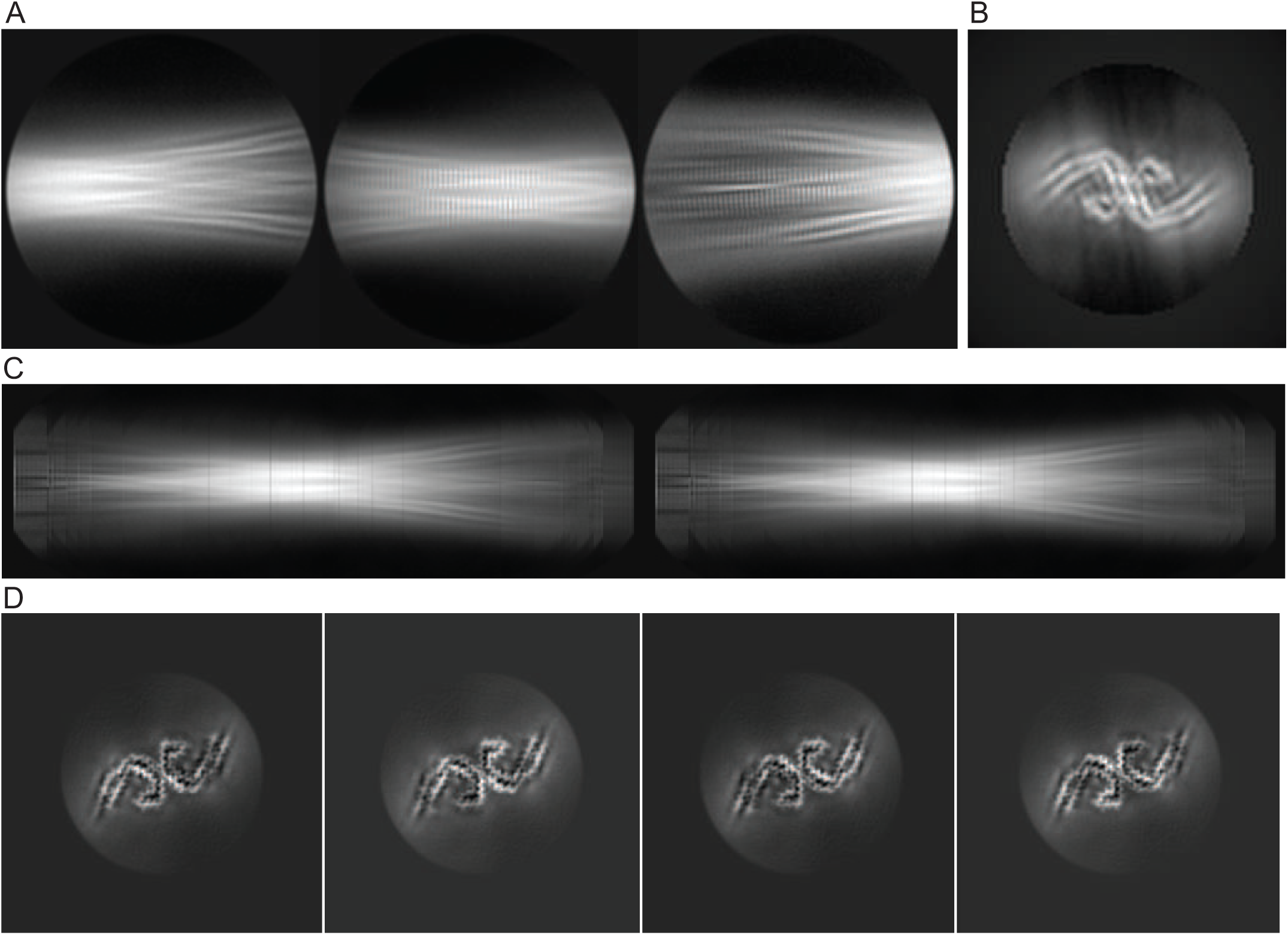
Initial model generation from multiple 2D class averages in smaller boxes. **A.** Three representatives of the twenty four 2D class averages that were used for initial model generation. **B.** The 2D reconstruction obtained from (half of) the pixel columns in **C. C.** The sinogram covering 360 degrees of rotation along the helical axis, as output by RELION. **D.** Four central slices of the symmetrised, post-processing map after two rounds of 3D auto-refinement, which started from the initial model that was generated from the image in B.

## 4. Discussion

Comparison of the maps obtained in sections 3.2 and 3.3 with the published 3.0 Å map of the EMPIAR entry (Figure 4) reveals that the map obtained in section 3.2 represents a local optimum. In particular, the relative heights of the different *β*-strands along the helical axis are incorrect, and the position and direction of some of the *β*-strands in the XY-direction are also incorrect. Therefore, despite its reasonable overall appearance and refinement statistics, this map could potentially have led to an incorrect tracing of the main-chain. The map obtained in section 3.3 closely resembles the published map. The small difference in resolution may be explained by the absence of particle polishing and further 3D classification, which were performed for the published map.

**Figure 4.**
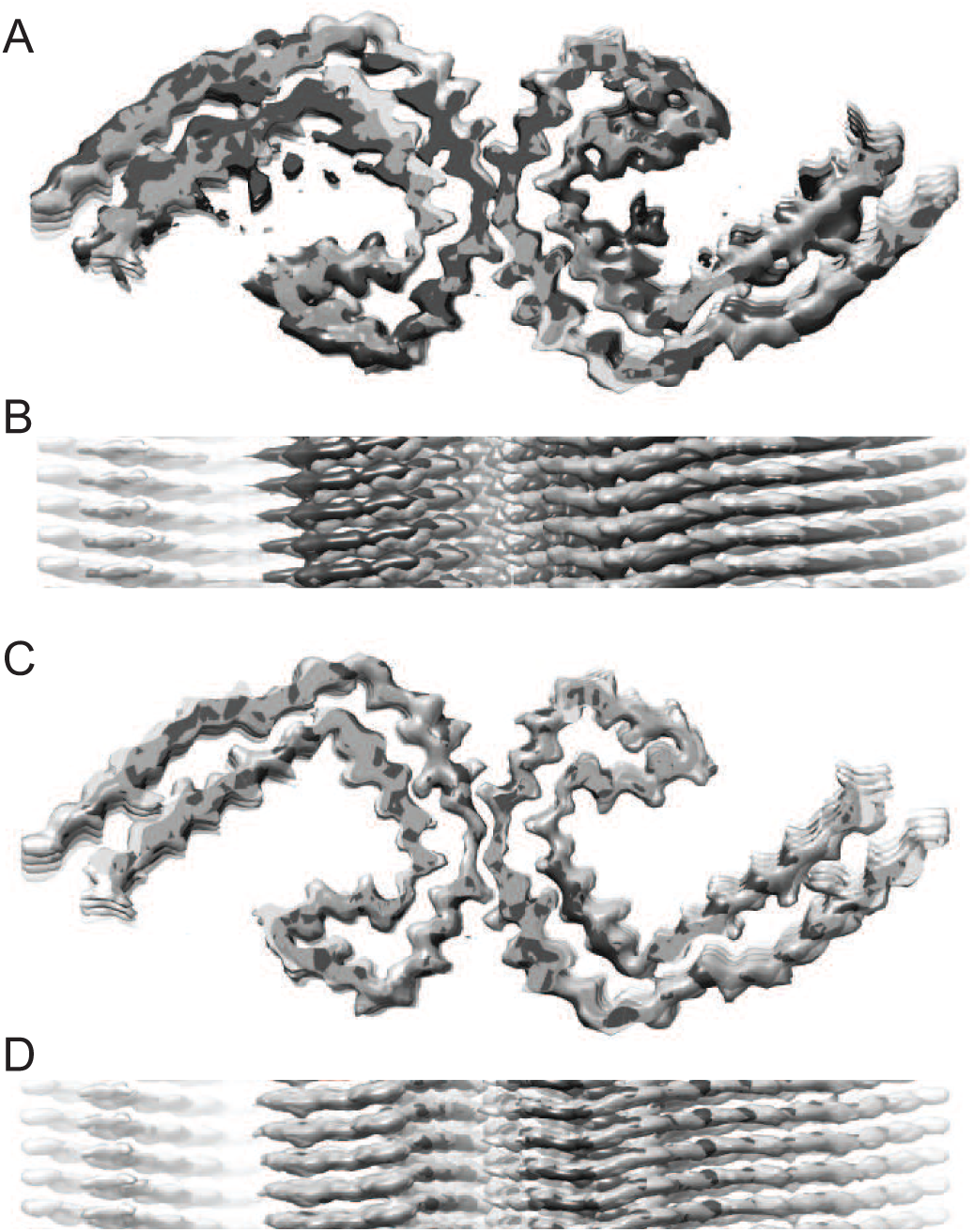
Comparisons with the published map. **A.** Overlay of the published map (transparent light grey) and the map after refinement of the initial model that was generated from a single 2D class average that spanned an entire cross-over (dark grey), in a view along the helical axis. **B.** As in A, but in a view perpendicular to the helical axis. **C.** As in A, but overlaying the published map (transparent light grey) and the map after two rounds of refinement, which started from the initial model that was generated from multiple aligned 2D class averages in smaller boxes. **D.** As in C, but in a view perpendicular to the helical axis.

The initial model that was generated from the 2D class averages with smaller boxes has higher resolution features than the model generated from the single 2D class average that spanned an entire cross-over. Yet, the resolution gain only happens in the XY direction, while at least part of the local optima of refinement seem to be related to the 4.75 Å signal in the Z direction. The proposed method of 2D reconstruction from 1D pixel columns is fast, which conveniently allows many trials to be performed in a short period of time. The method could in principle be extended to 3D reconstruction from 2D images, which would be slower. Meaningful signal in the Z-direction could only be obtained if the individual 2D class averages had sufficient detail to allow their accurate alignment along the helical axis, which is often not the case.

Different refinement strategies were used to obtain the maps described in sections 3.2 and 3.3. The former employed a single refinement, while the latter used two consecutive refinements. In addition, during the second refinement pseudo 2_1_ helical symmetry was imposed and the helical rise and twist were optimised. Performing the same strategy on the map obtained in section 3.2 also led to a correct solution (results not shown). Therefore, it seems that, for this data set, local minima can be avoided through the use of two consecutive refinements. This is not necessarily the case for all data sets. In fact, in our work on tau filament reconstructions, we have come across data sets of inferior quality, for which 3D refinements converged onto local optima more often, and these local optima persisted even after multiple rounds of refinements (results not shown).

Performing consecutive refinements with increasing initial resolutions comes with its own risk. In the second refinement in section 3.3, the initial resolution limit was set to 4.5 Å. This results in an initial reference in which *β*-strands are clearly separated, thereby allowing refinement of the helical twist and rise during all iterations of refinement. The clear separation of *β*-strands in the initial reference also prevents the two half-maps from converging onto different relative positions along the helical axis, as had happened in the first refinement in section 3.3. However, by using the combined half-maps from a previous refinement, the so-called gold-standard separation of half-sets is compromised, and the procedure becomes prone to overfitting of Fourier components with spatial frequencies up to the initial resolution limit. Therefore, it is important that the reported resolution of each refinement extends well beyond its initial resolution limit. In the example shown in this paper, the second 3D refinement was started at 4.5 Å resolution and converged onto a map with a resolution of 3.2 Å.

One problem that remains even when refinement has converged onto the global optimum is the determination of its absolute hand. For protein structures that contain *α*-helices, determination of the handedness is straighforward from the pitch of the helices, which becomes visible at resolutions beyond 5 Å. Because amyloids contain only *β*-strands, the handedness is often less clear. When resolution extends beyond 2.7 Å, the backbone oxygen atoms become visible, which again determines the handedness of the map. Due to the natural twist of *β*-strands, most amyloid filaments are left-handed, i.e. they have a negative helical twist angle in RELION. This however is not guaranteed, as cross-*β* packing interactions could in principle counteract the natural twist of *β*-strands. Two examples of such right-handed filaments are the ones formed by amyloid protein A in systemic amyloidosis (Liberta *et al.*, 2019), and by amyloid-*β* from the meninges in Alzheimer’s diseases (Kollmer *et al.*, 2019). Therefore, for resolutions below 2.7 Å, the handedness of the reconstruction may need to be inferred from the handedness of previously observed structures of homologous peptides, or may need to be determined using additional experiments, such as rotary shadowing (Stark *et al.*, 1984). The handedness of our tau PHF filament structures, which we had assumed to be left-handed based on the natural twist of *β*-strands, agreed with the handedness of tau filaments that were extracted from the brains of individuals with chronic traumatic encephalopathy, one of which was resolved to 2.3 Å resolution, showing clear densities for the backbone oxygens.

## 5. Conclusion

Refinement of amyloid structures may suffer from local optima that can lead to incorrect reconstructions with seemingly decent statistics. Densities with excellent local features in the XY-plane, but not in the Z-direction, or *vice versa*; variations in the intensity of protein densities; and negative densities in between the *β*-strands are signs that refinement may have converged onto a local optimum. A new method presented here uses reference-free 2D class averages to calculate initial 3D models that may be relatively close to the global optimum in the XY direction, but lack features along the helical axis. Even using these models, refinement of amyloid structures remains difficult, in particular for data with high noise levels. Therefore, *de novo* atomic modeling of amyloids should not be performed in RELION reconstructions with resolutions worse than 4 Å; better data should be collected instead. Maps with resolutions beyond 3.5 Å resolution, in which continuous main-chain density with convincing side chains are resolved in all three directions may provide increased confidence in the results.

I am grateful to Takanori Nakane for providing an image of his pet chicken; to Wenjuan Zhang, Yang Shi, Alexey Murzin, Takanori Nakane, Ben Falcon and Tony Crowther for discussions; and to Jake Grimmett and Toby Darling for help with high-performance computing. This work was supported by the EM-facility at MRC-LMB. Funding was provided by the UK Medical Research Council (MC UP A025 1013).

